## Supplementary Information for "Nanobody-based conjugates targeting small molecule-binding GPCRs and exhibiting logic-gated signaling"

| Compounds | $[M+H]^+_{calc}$ | $[M+H]^+_{obs}$ |
|---|---|---|
| CGS-azide | 699 | 700 |
| CGS-alkyne | 712 | 713 |
| 6E-azide | 1739 | 1742 |
| CGS-6E | 2451 | 2454 |
| $PTH_{1-11}$-Cys | 1412 | 1412 |
| $PTH_{1-11}$-DBCO | 1839 | 1839 |
| DynA8-Cys | 1084 | 1085 |
| DynA8-DBCO | 1513 | 1513 |
| G3-$PEG_4$-DBCO | 966 | 967 |
| G3-$PEG_{24}$-DBCO | 1847 | 1850 |
| G3-LL-$PEG_{24}$-DBCO | 2518 | 2506 |

**Supplementary Table 1: Mass spectrometry characterization of synthetic compound building blocks used in this study**. Compounds were analyzed by electrospray ionization mass spectrometry as described in Methods. Calculated masses ($[M+H]^+_{calc}$) refer to the monoisotopic mass of a singly protonated species. Masses recorded using mass spectrometry are labeled as $[M+H]^+_{obs}$. Sequences of peptides are shown **Supplementary Table 7**. Structures of small molecule building blocks are shown the Synthetic Methodology section.

| Conjugates | $[M+H]^+_{calc}$ | $[M+H]^+_{obs}$ |
|---|---|---|
| $Nb_{6E}$ | - | 13475 |
| $Nb_{6E}$-DBCO | 13239 | 13242 |
| CGS-$Nb_{6E}$ | 13939 | 13942 |
| $Nb_{ALFA}$ | - | 15208 |
| $Nb_{ALFA}$-DBCO | 14970 | 14977 |
| CGS-$Nb_{Alfa}$ | 15677 | 15678 |
| $Nb_{GFP}$ | - | 14235 |
| $Nb_{GFP}$-DBCO | 13999 | 14005 |
| CGS-$Nb_{GFP}$ | 14698 | 14693 |
| $Nb_{PTHR1}$ | - | 14467 |
| $Nb_{PTHR1}$-DBCO | 14231 | 14229 |
| CGS-$Nb_{PTHR1}$ | 14922 | 14932 |
| $Nb_{ALFA}$-biotin-azide | 14951 | 14944 |
| $Nb_{ALFA}$-$PEG_4$-$PTH_{1-11}$ | 17037 | 17037 |
| $Nb_{neg}$ | - | 14827 |
| $Nb_{neg}$-DBCO | 14591 | 14596 |
| CGS-$Nb_{neg}$ | 15291 | 15292 |
| $Nb_{MHC-I}$ | - | 14443 |
| $Nb_{MHC-I}$-DBCO | 14205 | 14205 |
| CGS-$Nb_{MHC-I}$ | 14905 | 14906 |
| CGS-$PEG_4$-$Nb_{PTHR1}$ | 15177 | 15178 |
| CGS-$PEG_{24}$-$Nb_{PTHR1}$ | 16061 | 16063 |
| CGS-$PEG_{24}$-$Nb_{PTHR1}$LL | 16704 | 16707 |
| $Nb_{PTHR1}$-biotin-azide | 14210 | 14207 |
| DynA8-$Nb_{PTHR1}$ | 15742 | 15718 |
| DynA8-$Nb_{neg}$ | 16109 | 16083 |
| $Nb_{GLP1R}$ | - | 14844 |
| $Nb_{GLP1R}$-DBCO | 14590 | 14610 |
| CGS-$Nb_{GLP1R}$ | 15307 | 15310 |

**Supplementary Table 2: Validation of Nb-ligand conjugate identity using mass spectrometry.** Nb-ligand conjugates were analyzed by mass spectrometry as described in Methods. $MW_{calc}$ refers to the calculated average molecular weight and $MW_{obs}$ refers to the molecular weight observed upon mass spectrometry followed by deconvolution as described in Methods. $Nb_{neg}$ corresponds to a nanobody targeting an unrelated GPCR (mGluR5), previously named Nb43,[53] which is not expressed in cells used in this study and serves as a negative control. Only experimentally observed masses are reported for unmodified nanobodies.

| $EC_{50}$ (±SEM), nM Ligands | peak cAMP | Washout AUC |
|---|---|---|
| **A2AR** | | |
| CGS | 2841 (896) | 1149 (281) |
| CGS-6E | >10000 | >10000 |
| CGS-$Nb_{6E}$ | >10000 | >10000 |
| CGS-$Nb_{ALFA}$ | >10000 | >10000 |
| CGS-$Nb_{GFP}$ | >10000 | >10000 |
| **A2AR-BC2-6E-ALFA** | | |
| CGS | 144 (25) | 714 (180) |
| CGS-$Nb_{6E}$ | 14 (4) | 7 (1.4) |
| CGS-$Nb_{ALFA}$ | 4.8 (1.3) | 5 (1) |
| CGS-$Nb_{GFP}$ | >10000 | >10000 |
| **A2AR-$Nb_{6E}$-ALFA** | | |
| CGS | 52 (8) | 95 (6.7) |
| CGS-6E | 1.8 (0.5) | 0.8 (0.2) |
| CGS-$Nb_{6E}$ | >10000 | >10000 |
| CGS-$Nb_{ALFA}$ | 2.6 (0.7) | 0.9 (0.3) |
| CGS-$Nb_{GFP}$ | >10000 | >10000 |
| **A2AR-GFP-ALFA** | | |
| CGS | 587 (26) | 2342 (768) |
| CGS-$Nb_{6E}$ | >10000 | >10000 |
| CGS-$Nb_{ALFA}$ | 31 (5) | 9 (2) |
| CGS-$Nb_{GFP}$ | 59 (32) | 23 (12) |

**Supplementary Table 3: Tabulation of the activity of ligands for inducing cAMP responses at A2AR and variants.** $EC_{50}$ values correspond to mean measurements from five independent experiments. Entries denoted “>10000” indicate that $EC_{50}$ values could not be determined due to weak ligand activity in that assay. “Peak cAMP” values are derived from fitting a sigmoidal concentration-response model to peak luminescence responses recorded after ligand addition (typically measured at 12 m). “Washout” data are derived from fitting a sigmoidal concentration-response model to integrated area under the curve values from sequential measurements performed every 2 m for a total of 30 m following ligand removal. $E_{max}$ values are similar across active compounds and are not listed for clarity. Corresponding concentration-response graphs are shown in **Figure 2** and **Supplementary Figures**. Note that a portion of this table is reproduced in **Table 1** in the main text.

| PTHR1 (µg) | A2AR (µg) | % maximal cAMP response | | | | | | | | |
|---|---|---|---|---|---|---|---|---|---|---|
| | | $PTH_{1-34}$ | | | $Nb_{PTHR1}$-CGS | | | CGS | | |
| | | N1 | N2 | N3 | N1 | N2 | N3 | N1 | N2 | N3 |
| 0.5 | 0 | 100 | 100 | 100 | 5 | 7 | 6 | 4 | 1 | 5 |
| 0 | 0.5 | 8 | 9 | 17 | 20 | 15 | 26 | 100 | 100 | 100 |
| 0.1 | 1 | 30 | 39 | 69 | 40 | 73 | 89 | 100 | 100 | 100 |
| 1 | 0.1 | 100 | 88 | 110 | 68 | 55 | 43 | 79 | 64 | 100 |
| 0.5 | 1 | 54 | 63 | 73 | 68 | 77 | 79 | 100 | 100 | 100 |
| 1 | 0.5 | 59 | 100 | 48 | 66 | 71 | 69 | 100 | 100 | 100 |
| 1 | 1 | 100 | 56 | 58 | 88 | 55 | 44 | 100 | 100 | 100 |

**Supplementary Table 4: Tabulation of cAMP (luminescence) responses induced at maximal concentrations of ligands upon variation of transfection conditions.** Cells transfected with varying amounts of A2AR and PTHR1 encoding plasmids were exposed to indicated concentrations of ligands. Peak cAMP luminescence responses were recorded and normalized to maximal responses recorded on a per experiment basis for each compound at the highest tested concentration. These data were used to generate the heatmap shown in **Figure 4**. Numerical values correspond to normalized mean cAMP/luminescence responses observed in three independent biological experiments.

| **Ligands** | **GLP1R-6E** | | **GLP1R-6E + A2AR** | |
|---|---|---|---|---|
| | **$EC_{50}$ (±SEM), nM** | **$E_{max}$ (±SEM), (% GLP-1 activation)#** | **$EC_{50}$ (±SEM), nM** | **$E_{max}$ (±SEM), (% GLP-1 activation)#** |
| CGS | >10000 | 5 (4) | 339 (145) | 32 (6) |
| GLP-1 | 4.1 (1.5) | 100 | 3.6 (0.4) | 100 |
| CGS-$Nb_{6E}$ | >10000 | 2 (1) | 23 (6) | 27 (4) |
| CGS-$Nb_{GLP1R}$ | >10000 | 2.5 (2.2) | 34 (19) | 27 (7) |
| CGS-$Nb_{neg}$ | >10000 | 2.4 (2) | >10000 | 11 (2) |

**Supplementary Table 5: Tabulation of ligand-induced cAMP (luminescence) responses in cells expressing GLP1R-6E and/or A2AR**. Cells stably expressing GLP1R-6E were either used directly or first transiently transfected to express A2AR in addition to GLP1R-6E. Data corresponds to mean $EC_{50}$ and $E_{max}$ values derived from at least three independent experiments (see representative example in **Figure 5**). $E_{max}$ values are normalized to the activity of an index ligand under control conditions, which result from fitting of a 3-parameter sigmoidal concentration-response model. Table entries of ">10000" indicate that $EC_{50}$ values could not be determined due to weak ligand activity in that assay.

| $EC_{50}$ (±SEM), nM Ligands | MOR-6E | MOR-6E + PTHR1 |
|---|---|---|
| **$G_{i2}$ TRUPATH** | | |
| DAMGO | 11 (4) | 18 (9) |
| DynA8 | 24 (4) | 29 (9) |
| DynA8-$Nb_{PTHR1}$ | 126 (19) | 6 (2) |
| DynA8-$Nb_{neg}$ | 134 (43) | 106 (37) |
| **β-arrestin recruitment** | | |
| DAMGO | 1069 (393) | 1505 (644) |
| DynA8 | 675 (103) | 914 (135) |
| $PTH_{1-34}$ | >10000 | 24 (6) |
| DynA8-$Nb_{PTHR1}$ | >10000 | 75 (11) |
| DynA8-$Nb_{neg}$ | >10000 | >10000 |

**Supplementary Table 6: Tabulation of the activity of ligands for inducing $G_{i2}$ activation or β-arrestin recruitment upon expression of MOR and PTHR1.** Data correspond to mean $EC_{50}$ values derived from three independent experiments (see **Figure 5**). $EC_{50}$ values result from fitting of a sigmoidal concentration-response model to data points. Entries marked ">10000" indicate that ligand activity was too weak to calculate an $EC_{50}$ value.

| Peptide | Sequence |
|---|---|
| 6E | QADQEAKELARQIS |
| $PTH_{1-34}$ | SVSEIQLMHNLGKHLNSMERVEWLRKKLQDVHNF |
| $PTH_{1-11}$ | AVUEIQLMHQRG |
| GLP-1 | HAEGTFTSDVSSYLEGQAAKEFIAWLVKGRGC |
| DynA8 | YGGFLRRIC |

**Supplementary Table 7:** Sequences of peptides used in this study. Peptide synthesis is described in Methods.

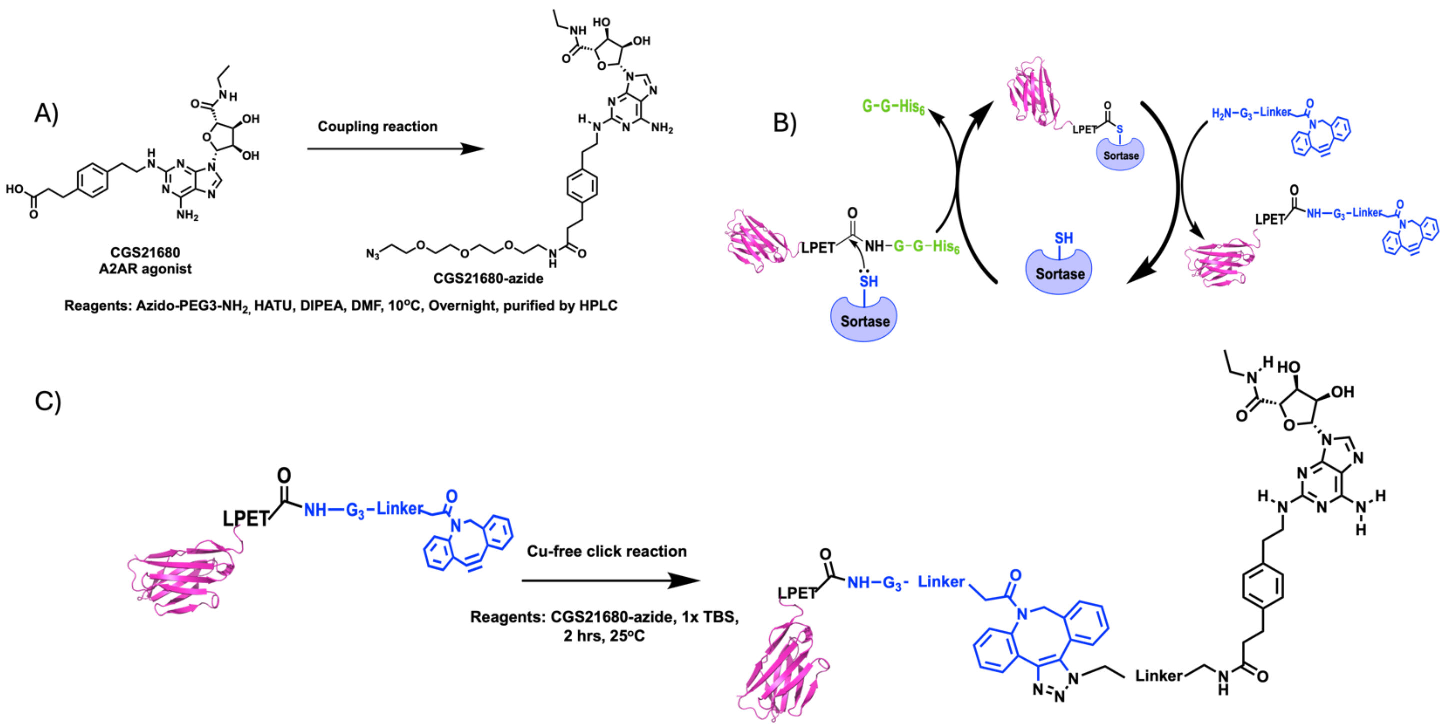


**Supplementary Figure 1: Synthesis of CGS-Nb conjugates used in this study. A**) Synthetic scheme for production of CGS-azide. Synthetic and purification details are descried in Methods. Mass spectrometry characterization of intermediates is found in **Supplementary Table 1**. **B**) Schematic representation of site-specific labeling of Nbs bearing a sortase recognition motif (LPETG) with DBCO using sortase A. Mass spectrometry characterization of Nb-DBCO conjugates used in this study can be found in **Supplementary Table 2**. **C**) Synthesis of CGS-Nb conjugates via copper-free azide-alkyne “click” chemistry. Detailed reaction conditions are provided in the synthetic methodology section of **Supplementary Methods**. Mass spectrometry characterization of Nb conjugates is shown in **Supplementary Table 2**.

A)

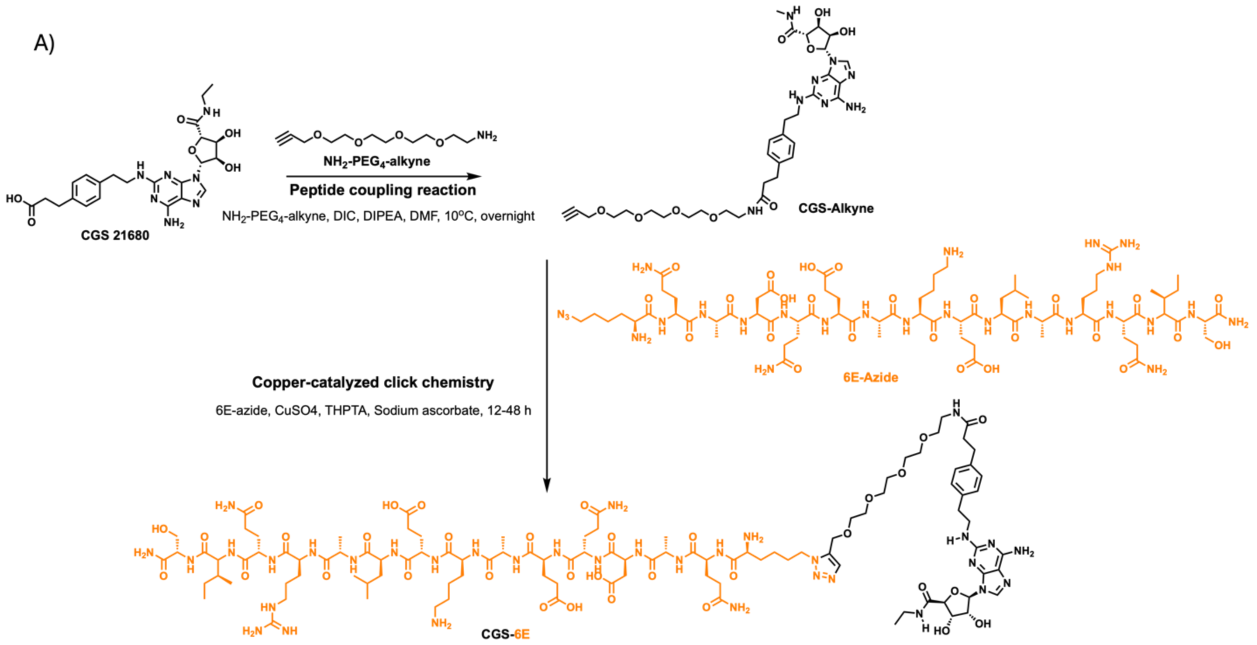


**Supplementary Figure 2: Synthesis of CGS-6E.** CGS-alkyne was synthesized as described previously[18] and in more detail in synthetic methodology section of **Supplementary Methods**.

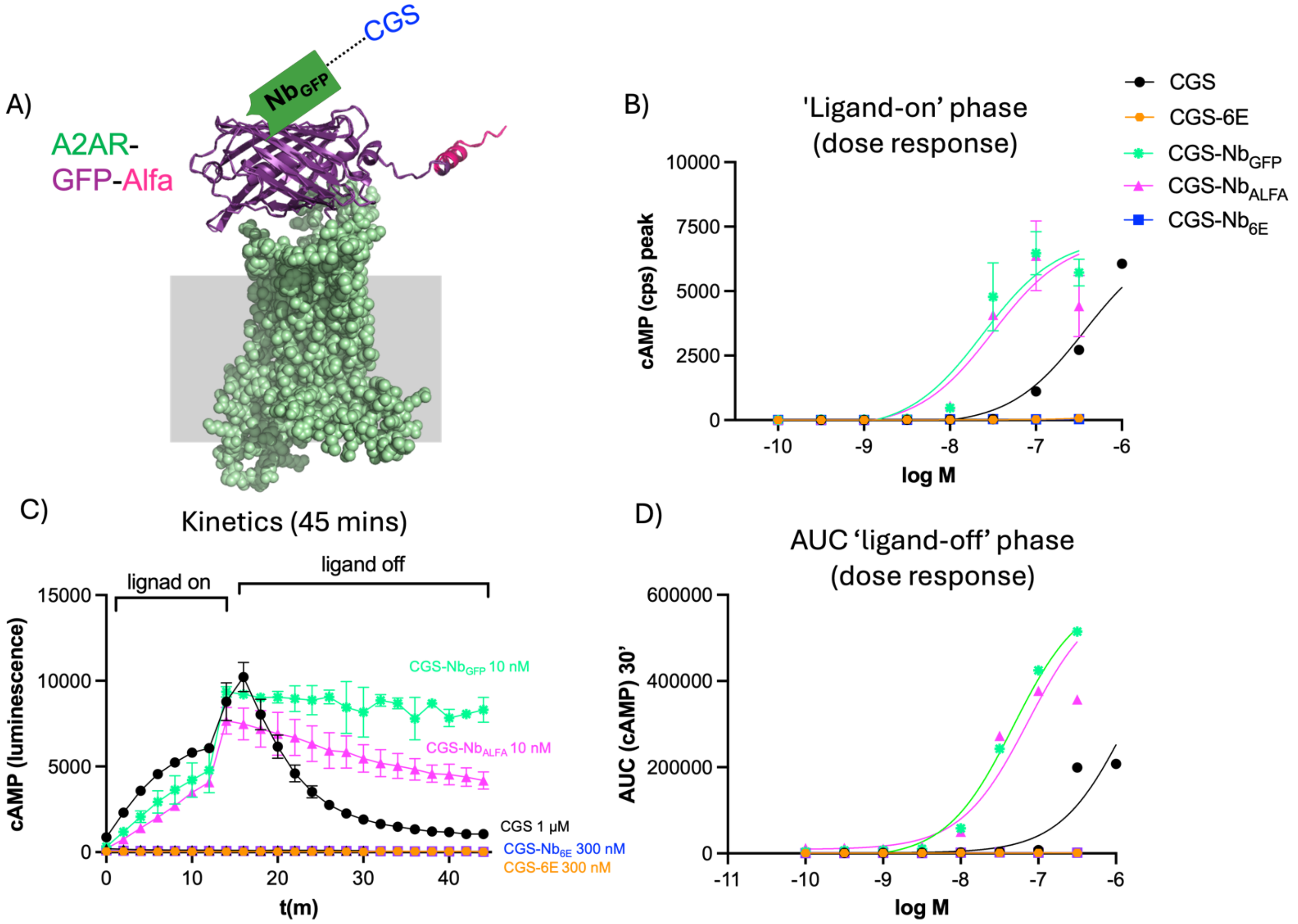


**Supplementary Figure 3: Assessment of ligand signaling at A2AR-GFP-ALFA. A**) Schematic of the receptor construct consisting of GFP and the ALFA epitope tag fused to the N-terminus (extracellular portion) of A2AR. This model of A2AR-GFP-ALFA was generated using Alphafold2 via Colabfold (see Methods). **B**) Representative concentration-response curves for ligand-induced cAMP production. Y-axis refers to the peak cAMP response signals recorded approximately 12 m after ligand addition. Responses are quantified by luminescence counts per second (cps). **C**) Representative kinetic measurements of ligand-induced cAMP responses upon ligand addition (12 m, "ligand on") and after ligand washout ("ligand-off). Responses were measured every 2 m. All data points correspond to mean ± S.D. from an individual representative experiment with two technical replicates. **D**) Washout responses assessed from quantifying area under curve of kinetic curves for "ligand off" portion. Data points correspond to mean ± standard error from technical duplicates in a representative experiment. Concentration-response curves were generated using a three-parameter logistic sigmoidal model. Summarized data that incorporates 3-5 biological replicate is shown in **Supplementary Table 3**.

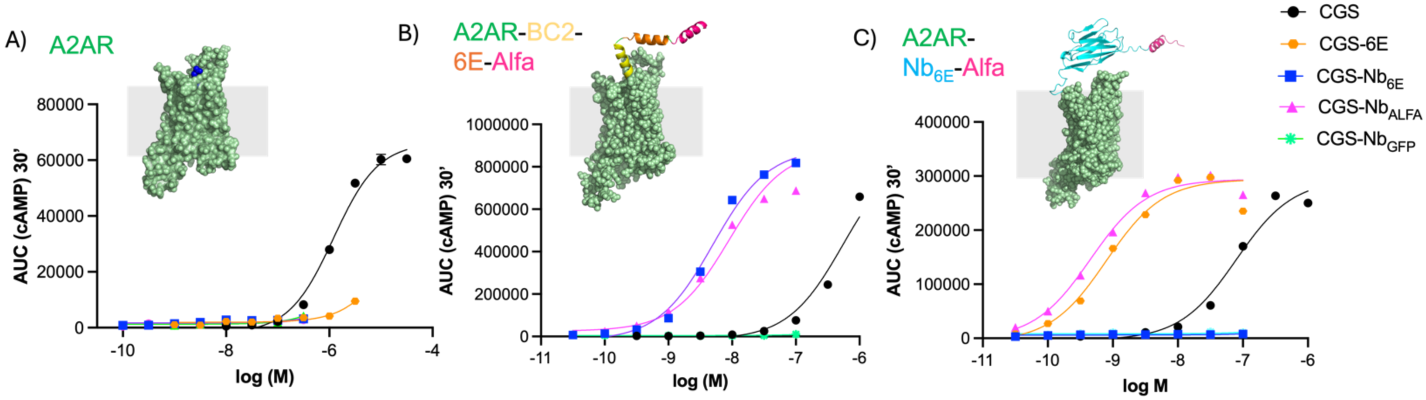


**Supplementary Figure 4: Measurement of A2AR cAMP signaling upon ligand washout.** Data points correspond to area under the curve integrations from kinetic measurements of cAMP responses upon ligand washout for 30 m for graphs shown in **Figure 2** for **A**) A2AR, **B**) A2AR-BC2-6E-ALFA, and **C**) A2AR-Nb6E-ALFA. Curves were generated by fitting data to 3-parameter sigmoidal dose-response models. Data correspond to mean ± standard error from an individual experiment with two technical replicates. Tabulated data that incorporates 3-5 biological replicate is shown in **Table 1.**

A)

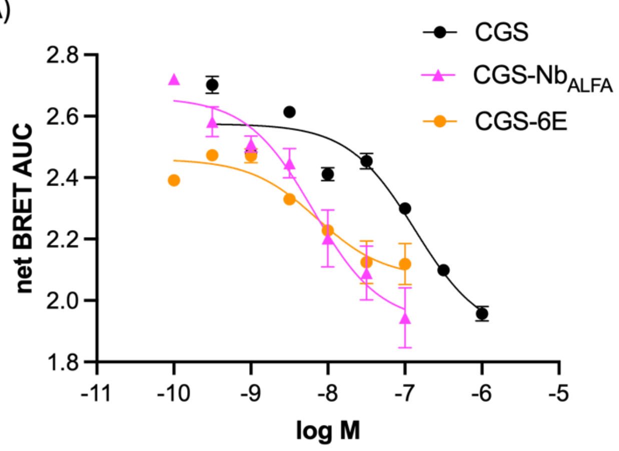


B)

| Ligands EC50 (± SEM), nM | Gαs-protein dissociation |
|---|---|
| CGS | 31 (17) |
| CGS-Nb$_{ALFA}$ | 5.4 (1.4) |
| CGS-6E | 5.7 (2) |

**Supplementary Figure 5: Characterization of ligand signaling at A2AR-Nb$_{6E}$-ALFA using a G-protein activation assay based on BRET. A**) The ligand-induced dissociation of Gαs protein was assessed in cells expressing A2AR-Nb$_{6E}$-ALFA. G-protein activation corresponds to a decrease in BRET between tagged G$_{\alpha s}$ and plasma membrane as described previously[30] (see Methods). Data corresponds to integrated area under the curve values generated from kinetic BRET measurements. Data points correspond to mean ± standard error from technical replicates in a single representative experiment. Curves are fit to a three-parameter logistic sigmoidal model. **B**) Tabulation of agonist potency parameters derived from 3 independent experiments. Data are presented as mean (±SEM).

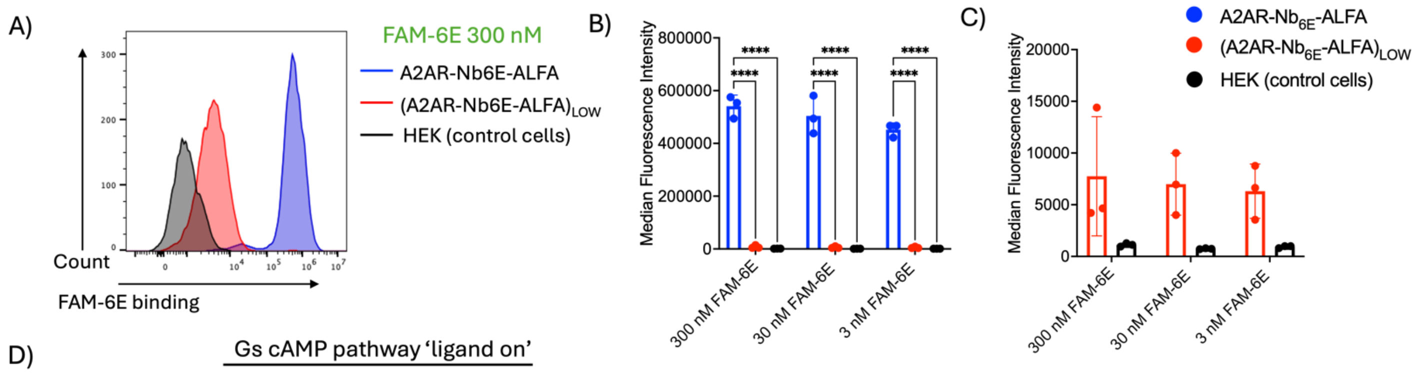


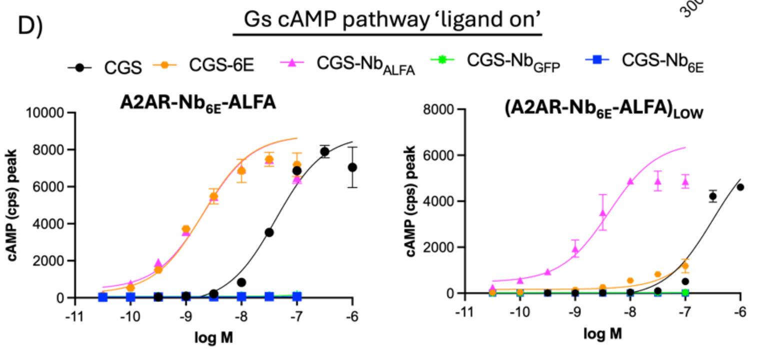


E)

| Ligands EC50 (± SEM), nM | cAMP pathway | |
|---|---|---|
| | A2AR-Nb6E-ALFA | (A2AR-Nb6E-ALFA)LOW |
| CGS | 52 (8) | 328 (89) |
| CGS-6E | 1.8 (0.5) | >10000 |
| CGS-Nb6E | >10000 | >10000 |
| CGS-NbALFA | 2.6 (0.7) | 5.4 (2.1) |
| CGS-NbGFP | >10000 | >10000 |

**Supplementary Figure 6: Evaluation of the impact of receptor expression levels on ligand signaling. A**) Analysis of the binding of fluorescein-labeled 6E peptide (described previously)[17] binding to clonal cell lines expressing either high (A2AR-Nb6E-ALFA, used throughout remainder of manuscript), or low (A2AR-Nb6E-ALFALOW) levels of receptor, and a negative control cell line not transfected with the receptor (HEK control cells). Cells were treated with FAM-6E (300 nM), followed by detection with a secondary (AF647-conjugated) anti-fluorescein antibody, as described in Methods. Live cells were gated on forward scatter/side scatter profiles and live cell staining is presented as a histogram of the APC channel. **B**) The intensity of cellular staining was quantified as median fluorescence intensity (MFI) values. Analysis of FAM-6E binding at various concentrations was performed with indicated cell lines. The bar graph data corresponds to mean ± SD from three independent experiments. Statistical significance was assessed by one-way ANOVA, with Dunnett's post hoc correction (*$p$ < 0.05; **p < 0.01; ***p < 0.001; ****$p$ < 0.0001; ns not significant). **C**) Expanded view of the data in the panel B excluding A2AR-Nb6E-ALFA to allow visualization of the differences in FAM-6E binding between (A2AR-Nb6E-ALFA)LOW and HEK control cells. These differences were not statistically significant. **D**) Representative concentration-response curves showing ligand-induced cAMP signaling in A2AR-Nb6E-ALFA and (A2AR-Nb6E-ALFA)LOW clonal cell lines. Y-axes refer to the peak cAMP response signals recorded approximately 12 m after ligand addition. Responses are quantified by luminescence counts per second (cps). **E**) Tabulation of agonist potency parameters derived from 3 independent experiments. These data incorporate the data shown in panel D. Table entries of ">10000" indicate that $EC_{50}$ values could not be determined due to weak ligand activity in that assay.

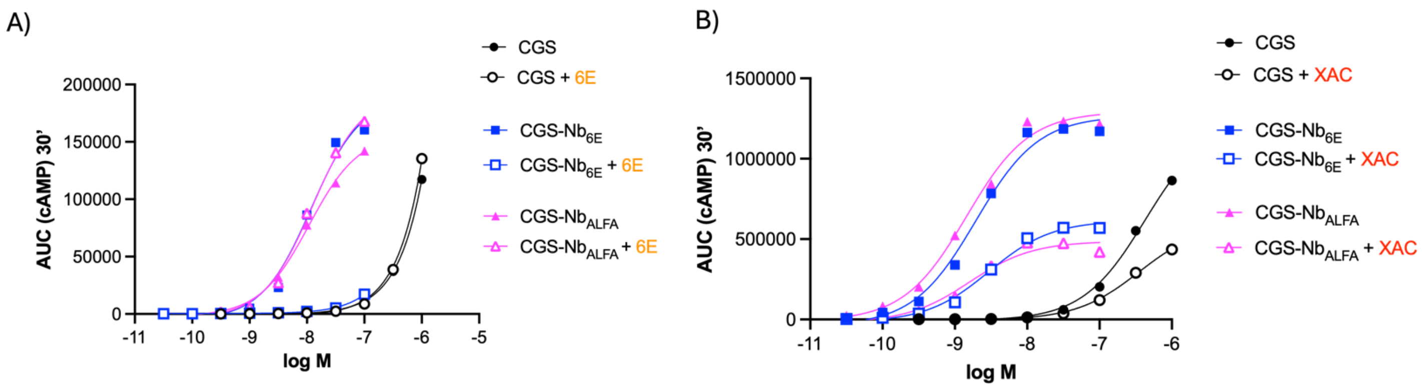


C)

| Ligands | cAMP, Washout AUC | | | | | |
|---|---|---|---|---|---|---|
| | Control | | +6E | | +XAC | |
| | EC50 (± SEM), nM | % CGS activation (± SEM)# | EC50 (± SEM), nM | % CGS activation (± SEM)# | EC50 (± SEM), nM | % CGS activation (± SEM)# |
| CGS | 578 (114) | 100 | 606 (129) | 115 (11) | 527 (186) | 76 (4) |
| CGS-$Nb_{ALFA}$ | 3 (1) | 100 (21) | 8 (3) | 110 (22) | 6.4 (1.2) | 53 (12) |
| CGS-$Nb_{6E}$ | 3.8 (1.4) | 110 (17) | 144 (81) | 30 (18) | 10.9 (3) | 68 (19) |

**Supplementary Figure 7: Impact of antagonists on ligand washout responses at epitope-tagged A2AR.** Concentration response curves were generated for cAMP washout signaling for ligands acting on A2AR receptor variant (A2AR-BC2-6E-ALFA) upon addition of **A**) synthetic 6E peptide or **B**) orthosteric A2AR antagonist (xanthine amine congener, XAC) during the washout phase. Data corresponds integrated area under the curve values from kinetic washout responses, as shown in the main **Figure 3** and described in Methods. Data points correspond to mean ± standard error from technical replicates. Data shown is from a representative experiment. **C**) Tabulation of agonist potency and $E_{max}$ (% CGS activation) parameters derived from 3 independent experiments performed as described in panels A-B. # Maximal cAMP response signals are normalized to the activity of an index ligand under control condition. Data are presented as mean (±SEM). Entries denoted as “>10000” indicate activity was too weak to calculate an $EC_{50}$ value.

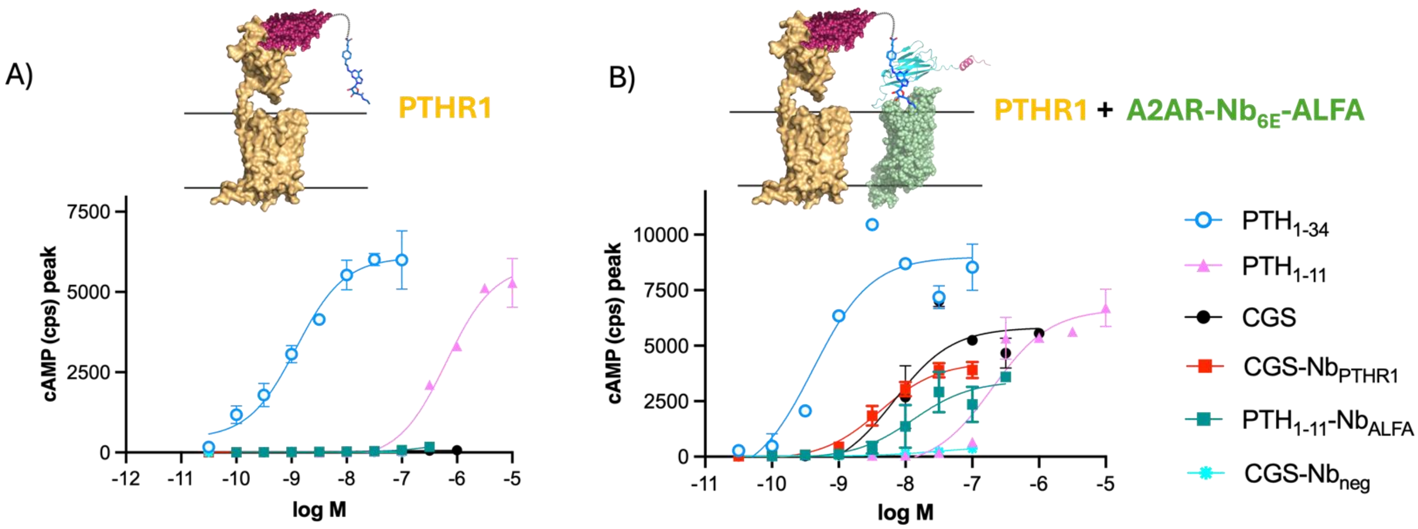


C)

| Ligands $EC_{50}$ (± SEM), nM | PTHR1 | PTHR1 + A2AR-$Nb_{6E}$-ALFA |
|---|---|---|
| $PTH_{1-34}$ | 22 (6) | 36 (16) |
| $PTH_{1-11}$ | 462 (31) | 492 (116) |
| CGS | >10000 | 238 (50) |
| CGS-$Nb_{PTHR1}$ | >10000 | 4 (1.4) |
| $PTH_{1-11}$-$Nb_{ALFA}$ | >10000 | 25 (7) |
| CGS-$Nb_{neg}$ | >10000 | >10000 |

**Supplementary Figure 8: Ligand signaling in cells stably expressing PTHR1 with transient expression of A2AR.** Representative concentration-response curves for cAMP induction in cells **A**) stably expressing PTHR1 (clonal cell line), and **B**) the same cell line (PTHR1) and transiently transfected to express A2AR-$Nb_{6E}$-ALFA. The schematic graphics depict the proposed mechanism of ligand activity in targeting receptor pairs. Unconjugated ligands (PTH1-34, PTH1-11, and CGS) were included as controls to validate receptor expression and functionality. Data corresponds to maximal cAMP responses recorded approximately 12 m after ligand addition and are quantified as luminescent counts per second (cps). Data points represent the mean ± SD from technical replicates in a representative experiment. Curves were fitted using a three-parameter logistic sigmoidal model. **C**) Tabulation of agonist potency parameters derived from 3-4 independent experiments that incorporates data shown in panels A and B. Table entries denoted ">10000" indicate that the $EC_{50}$ could not be determined due to weak ligand activity in that assay.

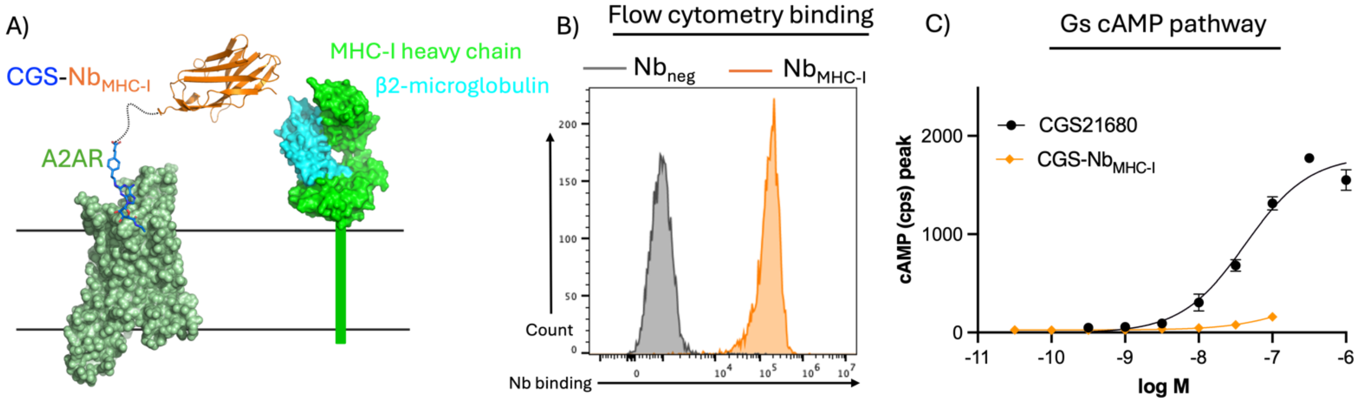


**Supplementary Figure 9: Attempted dual target engagement spanning A2AR and MHC-I. A**) Schematic representation of a CGS-Nb conjugate engaging a GPCR (A2AR) and MHC-I (non GPCR cell surface protein). **B**) Flow cytometry analysis of MHC-I expression on HEK293 cells through staining with a biotin-labeled Nb ($Nb_{MHC\text{-}I}$). $Nb_{neg}$ is a negative control nanobody (previously named Nb43)[53] that binds mGluR5 (not present on HEK293 cells). Cells were treated with 300 nM Nb-biotin, washed, and detected with streptavidin-APC, as described in Methods. Data is presented as a histogram of staining of live cells in the APC channel. **C**) Representative concentration-response graph for ligand-induced cAMP production a clonal cell line stably transfected to express A2AR receptor variant (A2AR-$Nb_{6E}$-ALFA) and endogenous expression of MHC-I. Maximal cAMP responses were recorded approximately 12 m after ligand addition and are quantified as luminescent counts per second (cps). Data points represent the mean ± SD from technical replicates of a representative experiment, with comparable results obtained in three independent experiments not shown. Curves were fitted using a three-parameter logistic sigmoidal model.

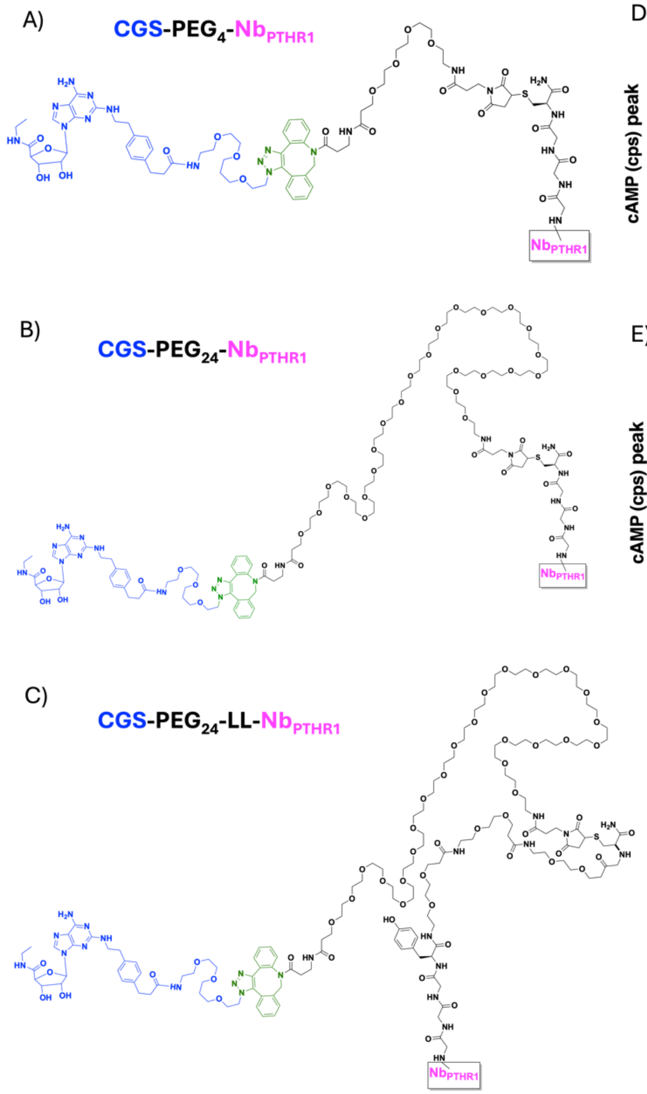


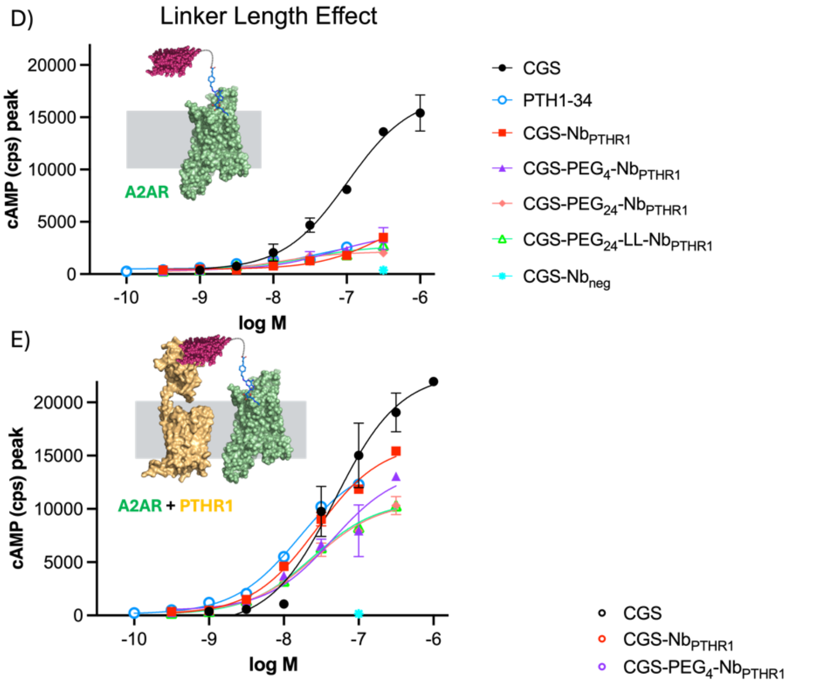


F)

| Ligands EC$_{50}$ (± SEM), nM | A2AR | A2AR + PTHR1 |
|---|---|---|
| CGS | 125 (26) | 92 (31) |
| PTH$_{1-34}$ | >10000 | 75 (57) |
| CGS-Nb$_{PTHR1}$ | >10000 | 33 (3) |
| CGS-PEG$_4$-Nb$_{PTHR1}$ | >10000 | 42 (9) |
| CGS-PEG$_{24}$-Nb$_{PTHR1}$ | >10000 | 27 (1.3) |
| CGS-PEG$_{24}$-LL-Nb$_{PTHR1}$ | >10000 | 33 (6) |
| CGS-Nb$_{neg}$ | >10000 | >10000 |

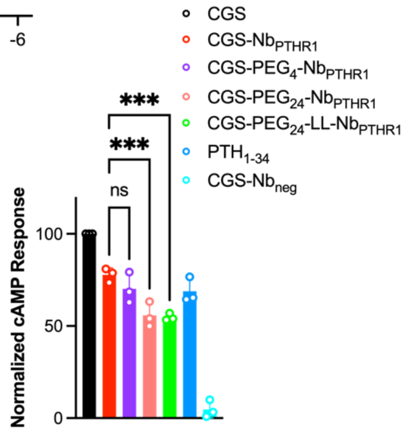


**Supplementary Figure 10: Evaluation of the contribution of linker length to CGS-conjugate activity. A-C**) Illustration of Nb-CGS conjugates with linkers of varying lengths. Linkers consist of linear polyethylene glycol (PEG) chains of **A**) 4- or **B**) 24-units, or **C**) an alternative linker comprised of a 24-PEG unit linker with an additional spacer, named PEG$_{24}$-LL. Compounds were synthesized as described in the synthetic methodology section of **Supplementary Methods**. Mass spectrometry characterization of the conjugates used in this study are provided in **Supplementary Tables 1 and 2**. **D-E**) Representative concentration-response curves for cAMP production in a HEK cells transiently transfected to express **D**) A2AR or **E**) A2AR and PTHR1. Data points correspond to mean ± SD from technical replicates in a single representative experiment (from three total biological replicates). Curves are fit to a three-parameter logistic sigmoidal model. **F**) Tabulation of agonist potency parameters derived from 3-4 independent experiments that incorporates data shown in panels C and D. Bar graph shows compilation of maximal luminescence (cAMP) responses produced by ligands at

A2AR at the highest concentrations tested. Luminescence responses observed at the highest ligand concentrations (300 nM for Nb-CGS conjugates, 1000 nM for CGS) were normalized to the response stimulated by the highest concentration of index ligand (CGS). Data in the bar graph show the mean ± SD from three independent experiments. Statistical significance was assessed by one-way ANOVA, with Dunnett's post hoc correction (**p* < 0.05; **p < 0.01; ***p < 0.001; *****p* < 0.0001; ns not significant).

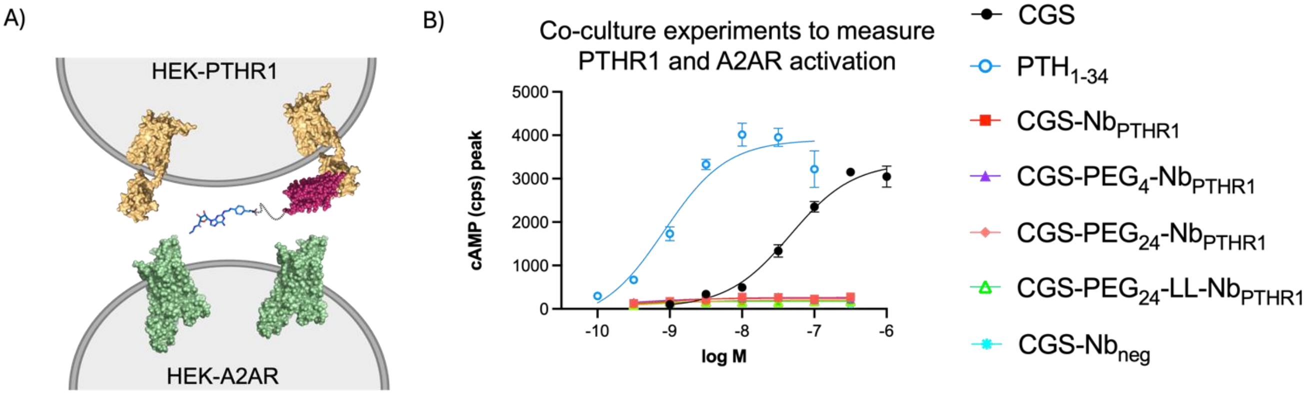


C)

| **Ligands** **$EC_{50}$ (± SEM), nM** | **Coculture PTHR1 and A2AR** |
|---|---|
| $PTH_{1-34}$ | 4 (3) |
| CGS | 52 (5) |
| CGS-$Nb_{PTHR1}$ | >10000 |
| CGS-$PEG_4$-$Nb_{PTHR1}$ | >10000 |
| CGS-$PEG_{24}$-$Nb_{PTHR1}$ | >10000 |
| CGS-$PEG_{24}$-LL-$Nb_{PTHR1}$ | >10000 |
| CGS-$Nb_{neg}$ | >10000 |

**Supplementary Figure 11: Assessment of GPCR activation by Nb-ligand conjugates in a co-culture context. A**) Schematic of the proposed mechanism of receptor activation across neighboring cells by Nb-ligand conjugates. **B**) Concentration-response curves for ligand-induced cAMP production in a mixed population of clonal cells expressing either PTHR1 or A2AR. Maximal cAMP responses were recorded approximately 12 m after ligand addition and are quantified as luminescent counts per second (cps). Data points represent the mean ± SD from technical replicates of a representative experiment. Curves were fitted using a three-parameter logistic sigmoidal model. **C**) Tabulation of agonist potency parameters derived from 3-5 biological replicates that incorporates data shown in panel B. Table entries denoted ">10000" indicate that the $EC_{50}$ could not be determined due to weak ligand activity in that assay.

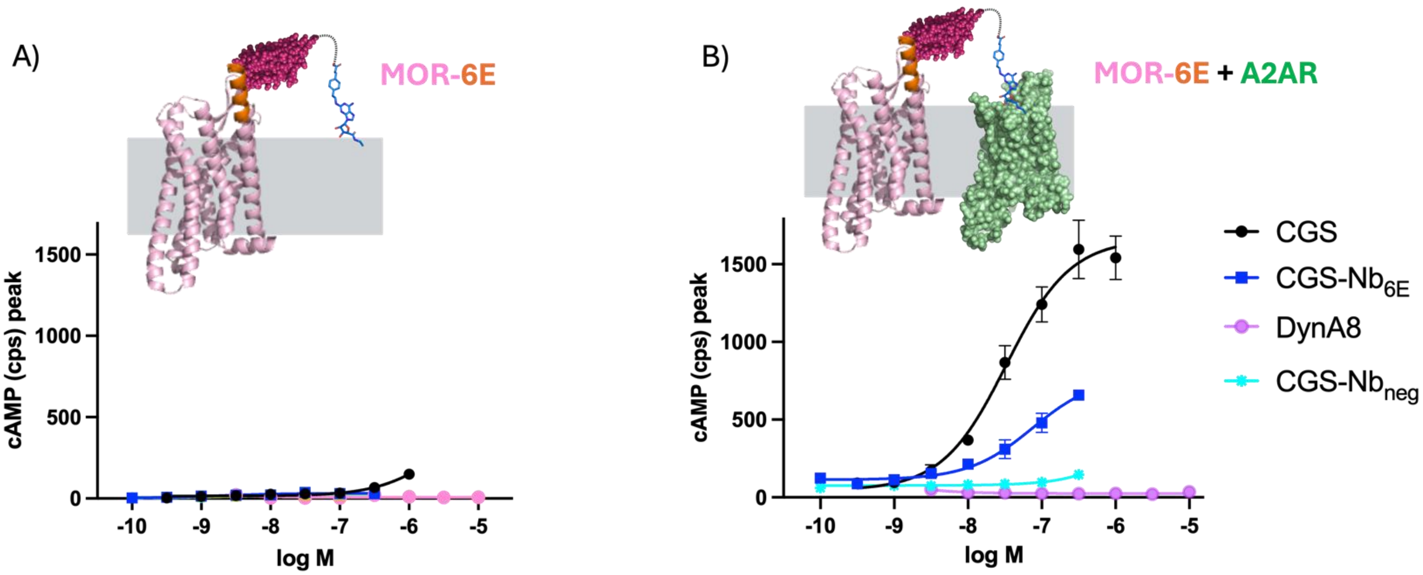


C)

| Ligands<br>EC50 (± SEM), nM | MOR-6E | MOR-6E + A2AR |
|---|---|---|
| CGS | >10000 | 29 (4) |
| CGS-$Nb_{6E}$ | >10000 | 43 (31) |
| Dyn A8 | >10000 | >10000 |
| CGS-$Nb_{neg}$ | >10000 | >10000 |

**Supplementary Figure 12: Extension of Nb-ligand conjugates to target alternate GPCR pairs.** Representative concentration-response curves for cAMP induction in a cell clonal cell line stably transfected to express **A**) MOR-6E, or **B**) that same cell line (MOR-6E) transiently transfected with A2AR. The schematic graphics depict the proposed mechanism of ligand activity in targeting receptor pairs. Maximal cAMP responses were recorded approximately 12 m after ligand addition and are quantified as luminescent counts per second (cps). Data points represent the mean ± SD from technical replicates in a representative experiment. Curves were fitted using a three-parameter logistic sigmoidal model. **C**) Tabulation of agonist potency parameters derived from 3 independent experiments that incorporates data shown in panels A and B. Table entries denoted “>10000” indicate that the $EC_{50}$ could not be determined due to weak ligand activity in that assay.

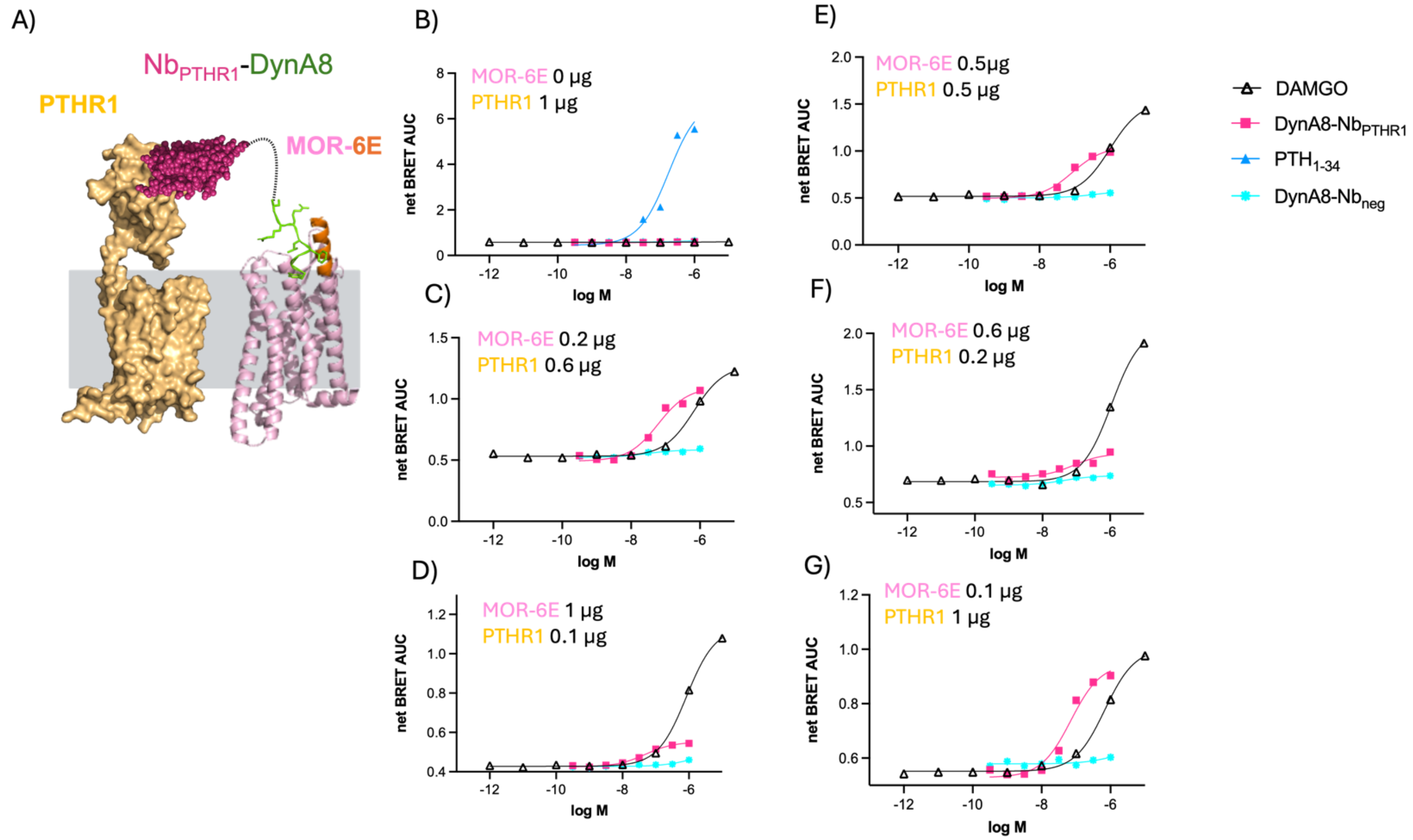


**Supplementary Figure 13: Assessment of the role of receptor expression levels on the induction of β-arrestin recruitment by Nb-ligand conjugates. A**) Schematic of the proposed mechanism of dual receptor targeting using Nb-ligand conjugates. **B-G**) Representative concentration-response curves for the induction of β-arrestin 2 recruitment in cells co-transfected with varying amounts of plasmids encoding A2AR and/or MOR-6E. Panel B includes PTH$_{1-34}$, which is excluded from subsequent panels for expanded view. β-arrestin 2 recruitment was measured using a BRET assay as described in Methods. Data points in panels B-G represent the mean ± SD from technical duplicates in a single representative experiment, with comparable results obtained in three independent experiments not shown. Curves correspond to fitting of data to a 3 parameter sigmoidal concentration-response model.

**Supplementary Methods:**

**Solid-phase peptide synthesis.**

Peptides were synthesized using solid phase peptide synthesis (SPPS) using Rink Amide ProTide resin (0.55 - 0.8 mmol/g), Fmoc protection of backbone amines, and standard

acid sensitive sidechain protecting groups, as previously described.[16,17] Synthesis was performed using a Liberty Blue 2 microwave peptide synthesis instrument (CEM). Where relevant, a Cys residue was incorporated at the C-terminus of the peptide for functionalization using Cys-maleimide chemistry (see below). N-terminal Fmoc deprotection was performed using 10% piperidine (v/v) in dimethylformamide (DMF). For coupling steps, Fmoc-protected amino acids (4 equivalents) were added with N,N-diisopropylcarbodiimide (DIC, 8 equivalents) and ethyl cyano(hydroxyimino)- acetate (Oxyma, 4 equivalents). Certain exotic amino acids, such as Fmoc-Lys($N_3$)-$CO_2$H (Chem-Impex, #29756) and Fmoc-Lys(biotin)-$CO_2$H (Chem-Impex, #04988) were added using manual coupling conditions. For these reactions, amino acid activation was performed with 7-Azabenzotriazol-1-yloxy)tripyrrolidinophosphonium hexafluorophosphate (PyAOP, AAPTech # CXZ070) and N,N-diisopropylethylamine (DIPEA).
Upon completion of peptide synthesis, the resin was washed with DMF, and then dichloromethane and dried via gentle air flow for 15 m. Peptides were cleaved from the resin using a mixture of trifluoroacetic acid (TFA), triisopropylsilane (TIPS), and $H_2O$ (95:2.5:2.5 by volume). After 2 h, TFA was evaporated using a gentle flow of nitrogen and peptides were precipitated in cold diethyl ether, followed by centrifugation (3000 g for 2 m) to pellet the crude precipitated peptide. The peptides were purified by reverse-phase high-performance liquid chromatography (RP-HPLC, see below). Mass spectrometry was used to confirm peptide identity, with characterization shown in **Supplementary Table 1**. Peptide sequences are shown in **Supplementary Table 7.**

**Reverse-phase high-performance liquid chromatography (RP-HPLC).**
Peptides were analyzed by reversed-phase HPLC using a preparative C18 column (Aeris PEPTIDE 5 μm XB-C18, LC Column 250 x 21.2 mm, AXIA packed) in a Shimadzu LC-20AR solvent delivery system at a flow rate of 10-15 mL/min using an eluent system of 0.1% (v/v) trifluoroacetic acid (TFA) in $H_2O$ and 0.1% TFA in acetonitrile (v/v) Fractions containing peptides of interest were identified using mass spectrometry analysis. Fractions of interest were combined and lyophilized. Lyophilized peptides are dissolved in DMSO at desired concentrations and stored at -20° C for further use.

**Mass spectrometry analysis.**
Mass spectrometry data were acquired on a Waters Xevo qTOF LC/MS. Samples were analyzed in positive ion mode. Data acquisition and processing were carried out using MassLynx software. Masses of large biomolecules were deconvoluted using the MaxENT function in MassLynx (Waters). The identities of each new preparation of synthetic compounds and their conjugates were characterized by mass spectrometry. Mass spectrometry characterization of ligands is provided in **Supplementary Table 1**.

**Nanobody expression and purification**
Nb protein sequences were codon optimized for bacterial expression and cloned into a pET26b expression in frame with pelB and His6 sequences using clone EZ service from GenScript. The expression plasmid was then transformed into E. coli BL21 (DE3) by heat shock. The transformed cells were cultured overnight at 37 °C with antibiotic (kanamycin) selection to prepare a starter culture. The next day, starter culture was used to inoculate a larger expression culture (1-4 L) containing kanamycin. This culture was grown at 37

°C until reaching an OD600 nm of 0.6−0.8. Protein expression was induced with isopropyl β-D-1-thiogalactopyranoside (IPTG) added to a final concentration of 1 mM and the culture was incubated overnight between 27 and 30 °C. The cells were harvested by centrifugation at 6000 rpm for 20 m (Avanti J Series centrifuge) and resuspended in running buffer (0.05 M Tris, 0.15 M NaCl, 0.02 M Imidazole; pH 7.5) containing protease inhibitor (Pierce Protease Inhibitor Tablets, Thermo Fisher A32953).
Resuspended cells were lysed on ice using ultra sonication via three cycles for 2 m at 30−40% power with 2 m cooling time between cycles. The lysate was centrifuged for 45 mins at 16,000 rpm to remove cell debris. The cleared supernatant was subjected to His-tag purification using batch-based Ni-NTA chromatography, and the bound protein was eluted with buffer containing imidazole (tris buffered saline + 150 mM imidazole, pH 7.5). The eluted sample was further purified by size-exclusion chromatography on a HiLoad 16/600 Superdex 200 pg column (Cytiva Akta Pure) using an isocratic TBS gradient at a flow rate of 1 mL/min. Fractions of interest were collected and concentrated using a 10 kDa molecular weight cutoff Amicon spin-concentrator. Protein identity was confirmed by mass spectrometry. Protein concentrations were determined with a Nanodrop spectrophotometer to measure absorption at 280 nm, which was converted to concentration using the protein extinction coefficient. The identity of all purified nanobodies were confirmed by mass spectrometry (**Supplementary Table 2**).

**Sortase-mediated Nb labeling reactions.**
Sortagging reactions were performed as previously described.[14] This reaction involves the following components: protein bearing a sortase recognition motif (LPETGG) followed by a $His_6$ tag at the C-terminus (20–200 μM final concentration), triglycine-probe conjugates (500–1000 μM final concentration), and Sortase 5M (30 μM final concentration). The reaction was carried out in Sortase buffer (50 mM Tris-HCl, 150 mM NaCl, 10 mM $CaCl_2$, pH 7.5) and incubated for 16 hours at 12 °C with agitation. Following the incubation, the reaction was incubated with nickel NTA beads to capture Sortase 5M and any unreacted starting protein. The uncaptured material was further purified using disposable desalting columns to remove triglycine-peptide probes (Cytiva PD-10 SephadexTM G-25M). Fractions containing the product were pooled and concentrated using spin filtration (Amicon Ultra 0.5 mL Centrifugal Filters 10 kDa NMWL). The molecular weight of the conjugates was confirmed by mass spectrometry and shown in **Supplementary Table 2**.

**Cell culture and transfection conditions.**
Cells were routinely cultured in Dulbecco's modified Eagle's medium (Gibco, Thermo Fisher Scientific) supplemented with 10% fetal bovine serum (Sigma-Aldrich) and 1x penicillin/streptomycin at 37 °C in 5% $CO_2$ and grown to 70−80% confluency in 75 $cm^2$ tissue culture flask before splitting. Human embryonic kidney 293 (HEK293) cells stably expressing a cAMP-responsive luciferase based pGlosensor-22F reporter were transfected with receptor plasmid using Lipofectamine 3000 Transfection Reagent (Thermo Fischer Scientific). Transfection was performed according to manufacturer instructions using Opti-MEM for dilution of transfection reagents (Thermo Fischer Scientific). Cells exposed to transfection reagent were incubated for 24 h under standard

culture conditions. All cell lines were regularly screened for mycoplasma infection using the Lonza MycoAlert mycoplasma detection kit and were found to be negative.

**Receptor constructs, generation of stable cell lines, and transient transfection conditions.**

The plasmids encoding A2AR, A2AR-Nb$_{6E}$-ALFA, PTHR1, PTHR1-6E, and GLP1R-6E have been previously described.[16–18] Two new cells lines that stably express either A2AR-6E-BC2-ALFA or MOR-6E were generated for this study. Sequence information for all receptor plasmids can be found below in **Supplementary Information**. Briefly, stable cell lines were generated by first transfecting the cell line stably expressing the cAMP-responsive luciferase variant with plasmids encoding receptors of interest and a geneticin resistance gene. Following transfection, cells were transferred into media containing the antibiotic G418 sulfate (0.5-1 mg/mL). Clonal cell lines were generated by limiting dilution to provide cell lines that stably expresses cAMP biosensor and receptor of interest without the need for continuous G418 selection. Transient transfections were conducted using a similar protocol; however, the cells were not subjected to antibiotic selection following transfection. Instead, they were directly transferred into a 96-well plate for subsequent assays.

**Flow cytometry assessment of cell staining.**

HEK293 cells stably expressing either native human A2AR or A2AR variants were cultured as described above. Cells were detached with trypsin and transferred to a round bottom 96 well plate and pelleted by centrifugation (500 rpm for 3 m). Cells were resuspended in phosphate-buffered saline (PBS) containing 2% bovine serum albumin (BSA) (w/v) (PBS/BSA). These cell preparations were incubated on ice with Nbs functionalized with biotin at concentrations ranging 0.1 pM – 1 μM for 30 min. Following incubation, cells were washed with PBS/BSA, centrifuged, and resuspended in PBS/BSA containing APC-conjugated streptavidin (1:2000 dilution) and incubated for 30 min on ice prior to washing. Washed cells were then resuspended in PBS/BSA for analysis by CytoFlex flow cytometer (Beckman Coulter). Live cells were gated based on forward scatter and side scatter profile and staining intensity was monitored in the APC channel. A minimum of 2,000 events corresponding to live cells were recorded. The data were further evaluated using the FlowJo 10 software.

**Synthetic methodology.**

*CGS-PEG3-azide.*

CGS-azide was prepared following previously described methods.[17] CGS21680 was dissolved in DMF along with 1 equivalent of 1-[Bis(dimethylamino)methylene]-1H-1,2,3-triazolo[4,5-b]pyridinium 3-oxide hexafluorophosphate (HATU) and stirred at room temperature for 30 minutes. Then 2 equivalents of $NH_2$-$PEG_3$-azide (Vector Laboratories, # CCT-AZ101) was added to the reaction mixture followed by addition of 5 equivalents of diisopropylethylamine (DIPEA). The reaction mixture was shaken at 10°C overnight. The reaction mixture was purified using reverse-phase preparatory HPLC and using a C18 column with a 20-70% gradient of acetonitrile in water containing 0.1% trifluoroacetic acid.

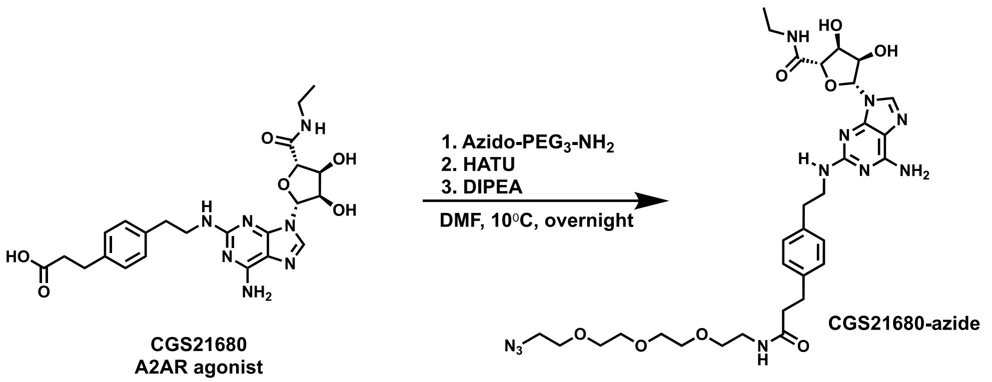


*CGS-6E*.
This compound was prepared as described previously (see **Supplementary Figure 2**).[17] CGS-alkyne was prepared using reaction conditions analogous to those used for the preparation of CGS-$PEG_3$-azide (see above). 6E-azide was synthesized using standard solid phase peptide synthesis methodology. CGS-6E was prepared by performing copper-catalyzed click chemistry between CGS-alkyne and 6E-azide. Briefly, 1 equivalent of 6E-azide was dissolved in DMF and 3 molar equivalents of CGS-alkyne was added to it. A 10x stock solution of $CuSO_4$ heptahydrate and THPTA was freshly prepared in water. This stock solution was used to add 20 equivalents of pre-mixed copper-THPTA solution to the solution of 6E-azide and CGS-alkyne. Lastly, 40 equivalents freshly prepared sodium ascorbate (from a 100 mM sodium ascorbate stock solution in water) was added. After overnight incubation, the product was purified using reverse-phase HPLC. Product identity was confirmed using mass spectrometry.

*GGG-DBCO*.
$G_3$-DBCO and other DBCO-labeled peptides were synthesized as previously described.[2,3] Briefly, GGG-Cys (100 mM; Genscript) and DBCO-maleimide (100 mM; VectorLabs # CCT-A108) were dissolved in dimethylsulfoxide (DMSO) supplemented with pH 7.4 phosphate buffer at a final concentration of 10 mM. After 4-5 h at 25°C, the reaction was purified using reverse-phase HPLC as described above. Other peptide-DBCO conjugates (DBCO-$PEG_4$-maleimide, Vector Laboratory, # CCT-A108P and DBCO-$PEG_{24}$-maleimide, BroadPharm # BP-25731) were prepared using analogous conditions.

*GGG-LL-Cysteine and G3LL-DBCO.*
GGG-LL-Cysteine was synthesized using the standard peptide synthesis protocol described above. Fmoc-NH-$PEG_2$-$CH_2CH_2$-COOH (ChemPep, # 280107) and Boc-$Gly_3$-OH (CHEM-IMPEX, # 07840) were used for synthesis. G3LL-DBCO was then synthesized using DBCO-$PEG_{24}$-maleimide using the conditions described above.

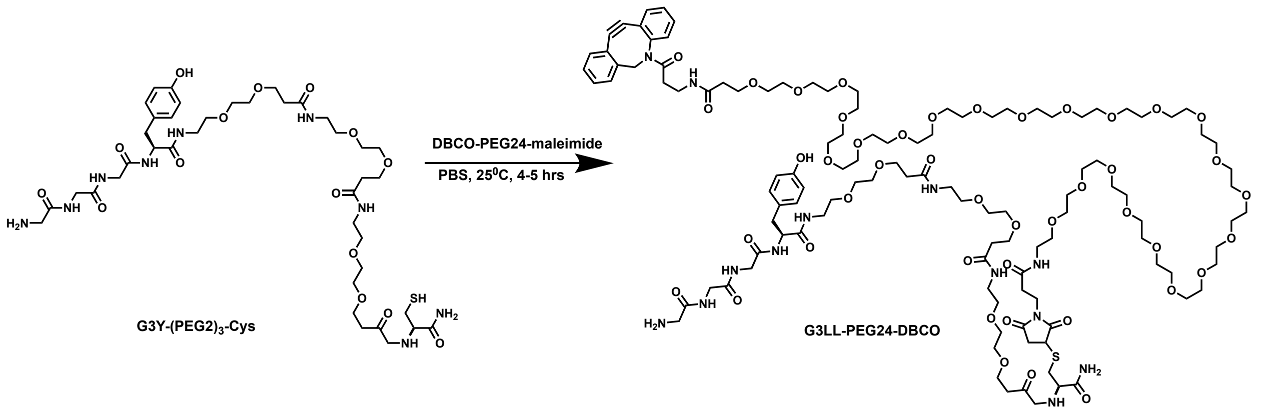


*Nanobody-small molecule (CGS) ligand conjugates.*
Nb-DBCO conjugates (produced using sortagging, described above) were mixed with 1.5 equivalents of CGS-PEG$_3$-azide in PBS buffer. The reaction proceeded for 2-3 h at which time the product (CGS-Nb conjugate) was purified using a Cytiva PD-10 SephadexTM G-25M size exclusion column to remove residual CGS-PEG$_3$-azide. The CGS-Nb conjugates are characterized by mass spectrometry (see **Supplementary Table 2**).

*Nanobody-peptide ligand conjugates.*
Here we describe the synthesis of DynA-Nb conjugates. Analogous methods were used for the synthesis of all Nb-peptide ligand (such as PTH-Nb) conjugates. DynA-DBCO was prepared using methods for Cys-maleimide reactions described above. Nb-biotin-azide conjugates were prepared using methods described in the sortagging methods section. Nb-biotin-azide was mixed with 3-6 equivalents of DynA-DBCO in PBS buffer. The reaction proceeded for 4 h at 25°C at which time product (DynA-Nb) purified using same size exclusion chromatography method as described above for CGS-Nb conjugates. The Nb-peptide ligand conjugates are characterized by mass spectrometry (see **Supplementary table 2**).

*Nanobody sequences.*
Nanobody sequences are taken from previously published work[14,16–18,35] or are listed below:

| Nanobody name: | Sequence or reference: |
|---|---|
| Nb$_{6E}$ | Cabalteja *et al.* ACS Chem Bio 2022 |
| Nb$_{ALFA}$ | Cabalteja *et al.* ACS Chem Bio 2022 |
| Nb$_{GFP}$ | Braga Emidio *et al.* Protein Science 2024 |
| Nb$_{PTHR1}$ | Sachdev *et al.* Nature Communications 2024 |
| Nb$_{GLP1R}$ | Sachdev *et al.* Nature Communications 2024 |
| Nb$_{mGluR5}$ (Nb$_{Neg}$ or previously Nb43) | QVQLVESGGGLVQAGGSLRLCAASGRTFTSYAMGWFRQA PGKERESVAAISSSGGSTHYADSVKGRFTISRDNSKNTVY LQMNSLKPEDTAVYYCAAAMYGSRWPDWEYDYWGQGTQVT VSSGGLPETGGHHHHHH |
| Nb$_{MHC-I}$ | Cheloha *et al.* RSC Chemical Biology 2021 |

Receptor sequences:

*A2AR-6E-BC2-ALFA:*

Amino acid:

MKTIIALSYIFCLVFAGPSRLEEELRRRLTEPGQADQEAKELARQISGPDRVRAVSHWSS
PIMGSSVYITVELAIAVLAILGNVLVCWAVWLNSNLQNVTNYFVVSLAAADIAVGVLAIPF
AITISTGFCAACHGCLFIACFVLVLTQSSIFSLLAIAIDRYIAIRIPLRYNGLVTGTRAKGIIAI
CWVLSFAIGLTPMLGWNNCGQPKEGKNHSQGCGEGQVACLFEDVVPMNYMVYFNFF
ACVLVPLLLMLGVYLRIFLAARRQLKQMESQPLPGERARSTLQKEVHAAKSLAIIVGLFA
LCWLPLHIINCFTFFCPDCSHAPLWLMYLAIVLSHTNSVVNPFIYAYRIREFRQTFRKIIRS
HVLRQQEPFKAAGTSARVLAAHGSDGEQVSLRLNGHPPGVWANGSAPHPERRPNGY
ALGLVSGGSAQESQGNTGLPDVELLSHELKGVCPEPPGLDDPLAQDGAGVSDYKDDD
DK*

Nucleotide:

atgaagaccatcatcgccctgagctacatcttctgcctggtgttcgccggccccagcagactggaagaggagctgagacg
gcgcctgacagagcctggacaggccgaccaggaggccaaggaactggctagacagatcagcggccctgatagagtg
cgggccgtgtcccactggtctagccccatcatgggctcctcggtgtacatcacggtggagctggccattgctgtgctggcca
tcctgggcaatgtgctggtgtgctgggccgtgtggctcaacagcaacctgcagaacgtcaccaactactttgtggtgtcact
ggcggcggccgacatcgcagtgggtgtgctcgccatcccctttgccatcaccatcagcaccgggttctgcgctgcctgcca
cggctgcctcttcattgcctgcttcgtcctggtcctcacgcagagctccatcttcagtctcctggccatcgccattgaccgctac
attgccatccgcatcccgctccggtacaatggcttggtgaccggcacgagggctaagggcatcattgccatctgctgggtg
ctgtcgtttgccatcggcctgactcccatgctaggttggaacaactgcggtcagccaaaggagggcaagaaccactccca
gggctgcggggagggccaagtggcctgtctctttgaggatgtggtccccatgaactacatggtgtacttcaacttctttgcctg
tgtgctggtgcccctgctgctcatgctgggtgtctatttgcggatcttcctggcggcgcgacgacagctgaagcagatggag
agccagcctctgccgggggagcgggcacggtccacactgcagaaggaggtccatgctgccaagtcactggccatcatt
gtggggctctttgccctctgctggctgcccctacacatcatcaactgcttcactttcttctgccccgactgcagccacgcccctc
tctggctcatgtacctggccatcgtcctctcccacaccaattcggttgtgaatcccttcatctacgcctaccgtatccgcgagtt
ccgccagaccttccgcaagatcattcgcagccacgtcctgaggcagcaagaacctttcaaggcagctggcaccagtgcc
cgggtcttggcagctcatggcagtgacggagagcaggtcagcctccgtctcaacggccacccgccaggagtgtgggcc
aacggcagtgctccccaccctgagcggaggcccaatggctatgccctggggctggtgagtggagggagtgcccaaga
gtcccaggggaacacgggcctcccagacgtggagctccttagccatgagctcaagggagtgtgcccagagcccctgg
cctagatgacccccctggcccaggatggagcaggagtgtccgattacaaggatgacgacgataagtga

*MOR-6E:*

Amino acid

MKTIIALSYIFCLVFAGQADQEAKELARQISGGDSSAAPTNASNCTDALAYSSCSPAPSP
GSWVNLSHLDGNLSDPCGPNRTDLGGRDSLCPPTGSPSMITAITIMALYSIVCVVGLFG
NFLVMYVIVRYTKMKTATNIYIFNLALADALATSTLPFQSVNYLMGTWPFGTILCKIVISID
YYNMFTSIFTLCTMSVDRYIAVCHPVKALDFRTPRNAKIINVCNWILSSAIGLPVMFMATT
KYRQGSIDCTLTFSHPTWYWENLLKICVFIFAFIMPVLIITVCYGLMILRLKSVRMLSGSK
EKDRNLRRITRMVLVVVAVFIVCWTPIHIYVIIKALVTIPETTFQTVSWHFCIALGYTNSCL
NPVLYAFLDENFKRCFREFCIPTSSNIEQQNSTRIRQNTRDHPSTANTVDRTNHQLENL
EAETAPLPDYKDDDDK

Nucleotide

atgaagaccatcatcgccctgagctacatcttctgcctggtgttcgccggccaggccgaccaggaggccaaggaactgg
ctagacagatctctggcggagacagcagcgctgcccccacgaacgccagcaattgcactgatgccttggcgtactcaag
ttgctccccagcacccagccccggttcctgggtcaacttgtcccacttagatggcaacctgtccgacccatgcggtccgaa
ccgcaccgacctgggcgggagagacagcctgtgccctccgaccggcagtccctccatgatcacggccatcacgatcat

ggccctctactccatcgtgtgcgtggtggggctcttcggaaacttcctggtcatgtatgtgattgtcagatacaccaagatgaa
gactgccaccaacatctacattttcaaccttgctctggcagatgccttagccaccagtaccctgcccttccagagtgtgaatt
acctaatgggaacatggccatttggaaccatcctttgcaagatagtgatctccatagattactataacatgttcaccagcatat
tcaccctctgcaccatgagtgttgatcgatacattgcagtctgccaccctgtcaaggccttagatttccgtactccccgaaatg
ccaaaattatcaatgtctgcaactggatcctctcttcagccattggtcttcctgtaatgttcatggctacaacaaaatacaggc
aaggttccatagattgtacactaacattctctcatccaacctggtactgggaaaacctgctgaagatctgtgttttcatcttcgc
cttcattatgccagtgctcatcattaccgtgtgctatggactgatgatcttgcgcctcaagagtgtccgcatgctctctggctcca
aagaaaaggacaggaatcttcgaaggatcaccaggatggtgctggtggtggtggctgtgttcatcgtctgctggactccca
ttcacatttacgtcatcattaaagccttggttacaatcccagaaactacgttccagactgtttcttggcacttctgcattgctctag
gttacacaaacagctgcctcaacccagtcctttatgcatttctggatgaaaacttcaaacgatgcttcagagagttctgtatcc
caacctcttccaacattgagcaacaaaactccactcgaattcgtcagaacactagagaccacccctccacggccaatac
agtggatagaactaatcatcagctagaaaatctggaagcagaaactgctccgttgcccgattacaaggatgacgacgat
aagtg
